# Chromosome-level Reference Genome Provides Insights into Divergence and Stress Adaptation of the African Oil Palm

**DOI:** 10.1101/2022.05.16.492201

**Authors:** Le Wang, May Lee, Zi Yi Wan, Bin Bai, Baoqing Ye, Yuzer Alfiko, Ramadsyah Ramadsyah, Sigit Purwantomo, Zhuojun Song, Antonius Suwanto, Gen Hua Yue

## Abstract

The palm family (Arecaceae), consisting of ∼ 2600 species, is the third most economically important family of plants. The African oil palm (*Elaeis guineensis*) is one of the most important palms. However, the genome sequences of palms available are still limited and fragmented. Here, we report a high-quality chromosome-level reference genome of an oil palm *Dura*. The genome of 1.7 Gb was assembled by integrating long reads with ∼ 150 × genome coverage. The assembled genome covered 94.5% of the estimated genome size, within which 91.6% were assigned into 16 pseudochromosomes and 73.7% were repetitive sequences. Relying on the conserved synteny with oil palm, the existing draft genome sequences of both date palm and coconut were further assembled into chromosomal level. Transposon burst, particularly long terminal repeat retrotransposons (LTRs) retrotransposons, following the last whole-genome duplication (WGD), likely explains genome size variation across palms. Convergent evolution of fruit colors tends to eliminate the roles of the *virescens* gene in controlling accumulation of anthocyanins in exocarp of ripe fruit of palms. Recent duplications of high tandemly repeated pathogenesis-related proteins (PRs) from the same tandem arrays played an important role in defense responses to *Ganoderma*. Whole genome re-sequencing of both ancestral African and introduced oil palms in Southeast Asia revealed that genes under putative selection were notably associated with stress responses, suggesting adaptation to stresses in the new habitat. The genomic resources and insights gained in this study could be exploited for accelerating genetic improvement and understanding the evolution of palms.

## Introduction

The palm family Arecaceae consists of ∼ 2600 species belonging to over 180 genera [1]. Over 90% of the diversity within this family is distributed in the tropical region of the world by adaptive radiation [2]. The Arecaceae is the third most economically important family of plants after the grasses and legumes [3]. The African oil palm (*Elaeis guineensis*) is of the most economic importance in Arecaceae, with a global production of ∼ 74 million metric tons of vegetable oil (FAOSTAT, http://www.fao.org/faostat, accessed at 2021/05/03). African oil palm is native to West Africa from Angola northward to Gambia [4]. It was introduced to Southeast Asia in 1840s, and has been naturalized since then [4]. Oil palm is the most productive oil plant and produces over 35% of vegetable oils with a market value of over $40 billion [5]. Although in the past 100 years, the oil yield has improved from ∼ 2.0 tons/ha/yr to the current ∼ 4.0 tons/ha/yr, there is still great potential to improve oil yield and other economical traits [4]. In addition, the oil palm industry is seriously threatened by diseases caused by *Ganoderma* species, resulting in losses of up to 80% of yield in some plantation areas [4]. Improvement of economically important traits using various approaches, including conventional and molecular breeding, is critically important in oil palm industry.

A high-quality genome assembly is necessary for both molecular breeding to accelerate genetic improvement and understanding species’ evolution. Despite the need to better understand oil palm genomics, only draft genome sequences are available. The completeness and quality of the published genome assembly are still to be improved [6-8]. Only ∼ 60% sequences out of the 1.8 Gb estimated genome were assembled and ∼ 45% sequences were anchored to genetic maps in *Pisifera* genome version EG5.1 and/or PMv6 [7, 8]. These draft genome sequences supply important tools to initiate molecular breeding to accelerate the genetic improvement. However, due to the limited completeness, fragmentation of scaffolds and incomplete annotations, their applications in genome-wide association studies, comparative genomics and structural variations in the species and its related species are limited. Therefore, further improvement of the draft genome of oil palm is essential for molecular breeding to improve economic traits and for comparative genomics to understand the evolution of palms [9].

Here, we report a high-quality chromosome-level genome sequence of *E. guineensis*. Comparative genomics revealed that transposon burst was responsible for genome size expansion in palms. We found evidences that high tandemly repeated pathogenesis-related proteins (PRs) played an important role in defense responses to *Ganoderma* infection. Whole-genome re-sequencing of 72 trees from West Africa and Southeast Asia revealed the population structure and lower genetic variations of oil palms in Southeast Asia. Signatures of local adaptation in the genome of oil palm was also found. The novel genomic resources and insights gained from this study will contribute to the understanding of evolution of palms and to accelerate the genetic improvement of oil palm.

## Results and discussion

### Chromosomal-level genome of African oil palm

Over 150 × coverage of long reads was assembled into 4752 contigs, with a total length of 1.7 Gb, covering 94.5% of the estimated genome (1.8 Gb) (Table S1). Contig N50 and the longest contig reached up to 2.168 and 12.851 Mb, respectively. We constructed five high-density linkage maps in five F_2_ populations, with the number of mapped markers ranging from 12,068 to 19,581 (Table S2; Figure S1). Anchoring contigs to these high-density genetic maps, based on a total number of 60,989 informative segregating markers, resulted in 16 pseudochromosomes consisting of 91.6% of assembled sequences and with a length ranging from 37.784 to 160.148 Mb, and 59.7% of assembled sequences were oriented (Tables S1, S3, and S4; Figure S2). Genome completeness analysis assessed by BUSCO showed that 95.8% of the core genes were found in the genome and 93.3 % were complete (Table S5). Mapping of assembled transcripts and *de novo* assembled RAD tags showed that 98.8% and 97.5% were matched to the genome assembly, respectively. We annotated long terminal repeats (LTRs). The LTR assembly index (LAI) was estimated to be 15.453 ± 2.968 (SD). This genome assembly significantly increases the total length of assembled sequences by ∼ 62%, N50 contig size of ∼233 folds, N50 scaffold size of ∼ 80 folds and total length of sequences anchored on pseudochromosomes of ∼ 2.4 folds, compared to previous draft genome sequences (Table S1) [6-8]. A chromosomal-level genome is necessary for comparative genomics to study genome duplications and understand the genomic architecture of adaptive radiation of palms. Date palm Barhee BC4 is one of the most impressive assemblies in palms, where < 50% of sequences were anchored to pseudochromosomes [10]. Although diverged at ∼ 65 million years ago (Mya) [7], we observed a high level of conserved chromosome synteny between oil palm and date palm (**Figure 1**), indicating that the chromosomal-level genome of oil palm can be used to for comparative genomic analysis. Taken together, our genome assembly showed high genome coverage, high assembly accuracy, long sequence continuity and high completeness of both genes and repetitive elements. Therefore, it will contribute to studies on genetics, genomics and breeding in palm species.

**Figure 1.**
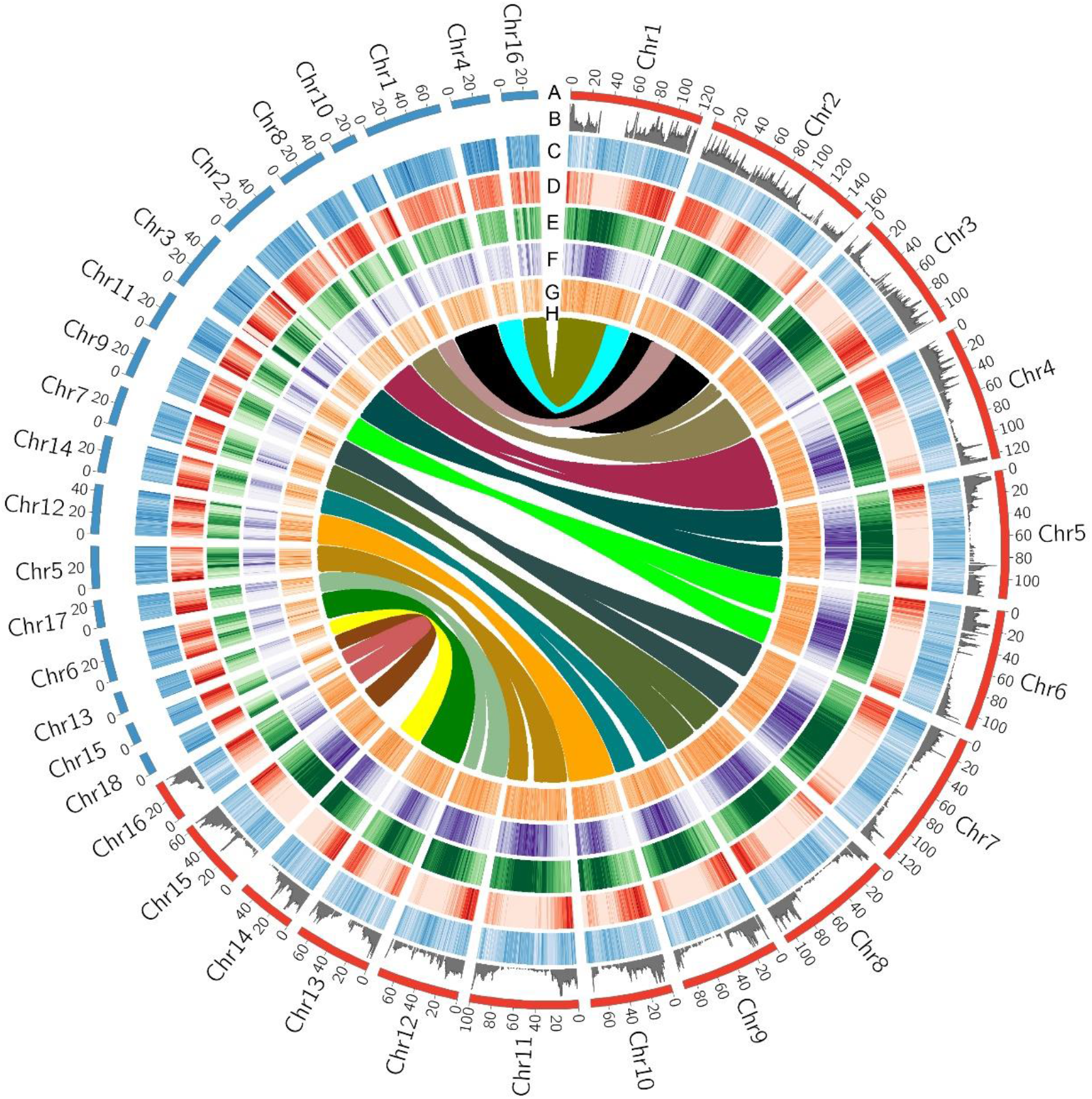
Global view of genomic features of oil palm and genomic synteny with date palm. **A**. Length of individual pseudochromosomes. **B**. Distribution pattern of recombination rate throughout individual chromosomes. **C**. Distribution pattern of GC content throughout individual chromosomes, estimated in 500-kb window. **D**. Distribution pattern of gene density throughout individual chromosomes, estimated in 500-kb window. **E**. Distribution pattern of repetitive sequences throughout individual chromosomes, estimated in 500-kb window. **F**. Distribution pattern of LTR retrotransposon superfamily Copia throughout individual chromosomes, estimated in 500-kb window. **G**. Distribution pattern of LTR retrotransposon superfamily Gypsy throughout individual chromosomes, estimated in 500-kb window. **H**. Conserved syntenic blocks between a pair of homologous chromosomes of oil palm and date palm. LTR, long terminal repeat.

### Annotations of the African oil palm genome

Repetitive sequences accounted for 73.7% of the genome assembly of oil palm (Table S6), significantly higher than previously observed both in the incomplete genome assembly of this species of ∼57% [7] and in date palm of ∼ 39 % [10]. In particular, LTRs took up 55.8 % of the genome. Copia is the largest class of LTRs, followed by the Gypsy superfamily, representing 39.5% and 17.2% of the assembled genome sequences, respectively (Table S7). The proportion of the two LTR superfamilies are also much higher than those in date palm of ∼ 14% and ∼ 4%, respectively [10]. We observed that the distribution pattern of repetitive sequences was negatively correlated to that of recombination rate (*R* = − 0.412, *P* < 10^− 4^) and gene density (*R* = − 0.794, *P* < 10^− 6^), but positively correlated to GC content (*R* = 0.932, *P* < 10^− 6^) (**Figure 1A**– **E**). In date palm, we observed the same pattern between repetitive sequences and gene density (*R* = − 0.856, *P* < 10^− 6^) and between repetitive sequences and GC content (*R* = 0.403, *P* < 10^− 6^) (**Figure 1A**–**E**) as in oil palm, which addresses how transposon dynamics have significantly shaped the genomic architecture of palms. We observed that distribution of Copia was highly correlated with that of the overall repetitive sequences (*R* = 0.952, *P* < 10^− 6^), while Gypsy were more likely randomly distributed across the genome (*R* = 0.107, *P* < 0.05) (**Figure 1E**–**G**). Our data indicates that palms have a much higher copy number of Copia than Gypsy, contradicting most other plant genomes that show higher Gypsy content [10]. Previous studies have shown retrotransposons in plants play important roles in genome size, genome structure remodeling, gene function, and genome evolution [11]. Therefore, it is highly possible that Copia may play an important role in the evolution of palms.

Gene annotations based on RNA sequencing, *ab initio* predictions, plant protein coding genes and protein domains, predicted 33,447 protein coding genes. Of these genes, 29,293 (87.6%) were annotated with known proteins or domains (Table S1). Over 95% of predicted genes showed an annotation edit distance (AED) value of < 0.5, indicating high-quality annotations of the genome (Figure S3). Median gene length was ∼ 5.2 kb, slightly higher than those of previous oil palm and date palm assemblies of ∼ 4.7 kb and ∼ 4.2 kb, respectively [7, 10]. In addition, more than 98.5% of the annotated genes were mapped to the 16 chromosome sequences, indicating that this genome assembly represents a nearly complete protein coding genome and is useful in future genetic and genomic studies. Functional enrichment analysis revealed that gene families showing expansions in oil palm were more involved in stress responses to pathogens and regulation of osmotic stresses (Figures S4 and S5).

### Transposon bursts lead to genome expansion and gene diversification in palms

The variation in genome size across eukaryotes is tremendous and is associated with species diversity [12]. Polyploidy and transposon expansion are the two major forces driving genome size variation, providing essential resources for evolutionary innovations, by generating novel genetic variations and altering gene expression patterns [13]. Thus, unraveling the mechanisms is necessary to understand the adaptive radiation and successful ecological dominance of the taxa. Genome size of palms varies from ∼ 800 Mb to ∼ 3Gb [14]. Oil palm and date palm show striking difference in genome size, with the predicted size of 1.8 Gb and 800 Mb, respectively, providing an excellent system to study genome size variation. Monocots share a common WGD event at ∼ 150 Mya [15]. The other paleopolyploid event, exclusively for the ancestor of all palms, occurred at ∼ 75 Mya, resulting in palaeotetraploidy of all palms [7, 14]. We observed large conserved syntenic blocks between homologous chromosome pairs throughout the whole genome (Figure S6), allowing examination of the effects of WGD events on genome evolution. Distribution of Ks, estimated based on 4,292 and 2,793 pairs of homologous genes from syntenic blocks in oil palm and date palm, respectively, revealed a major peak at ∼ 0.32, corresponding to the recent WGD at ∼ 75 Mya, shared by all palms (**Figure 2A** and **B**) [14]. A more recent Ks peak was observed at ∼ 0.22 for orthologous gene pairs, indicating the divergence between oil palm and date palm at ∼ 65 Mya [7]. Here, the divergence of the whole genome-wide homologous genes supports the conclusion that all palms have experienced two WGD events before adaptive radiation [14].

**Figure 2.**
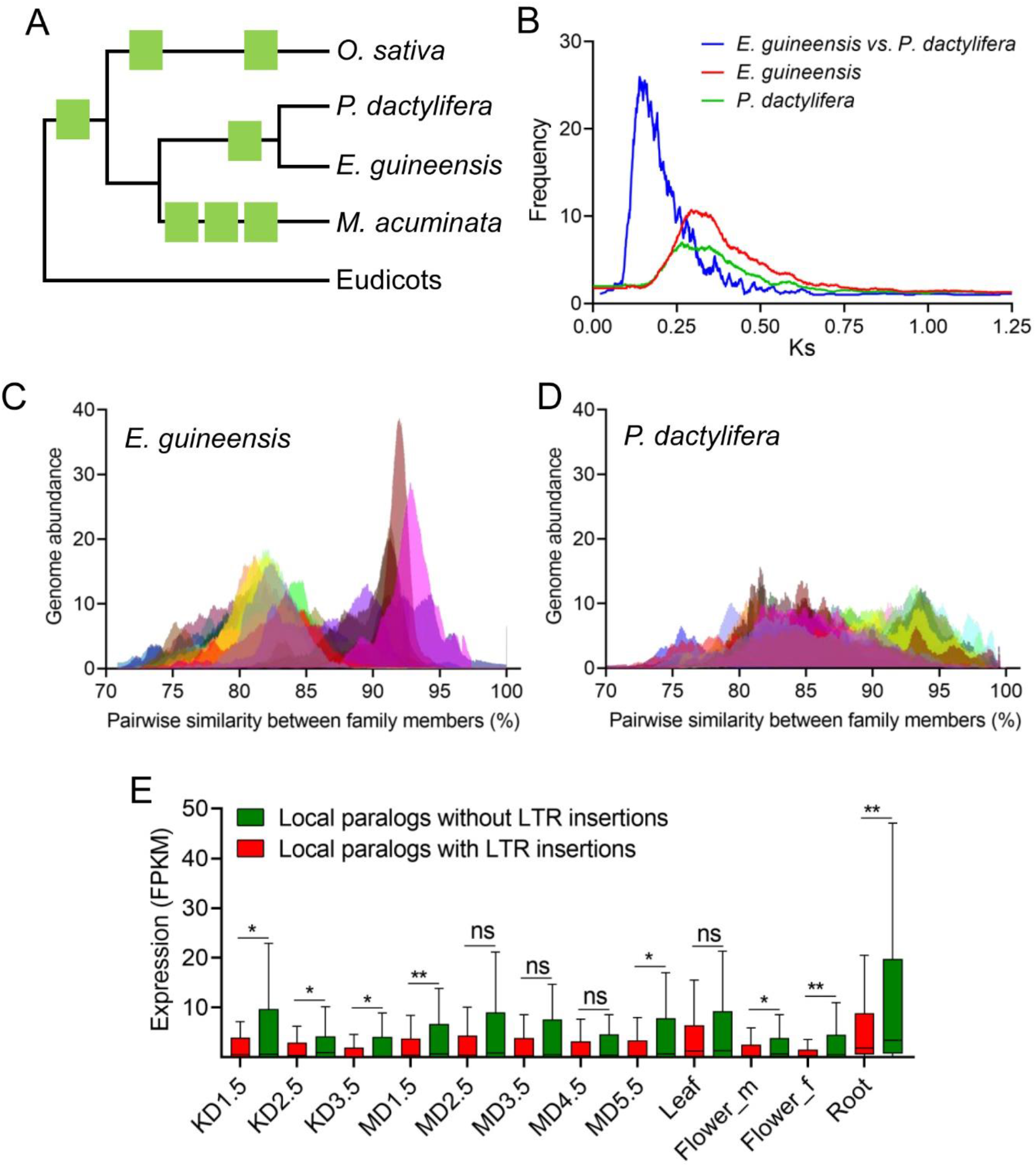
Transposon expansion drives genome evolution of palms. **A**. A phylogram showing historical whole-genome duplication (WGD) events in palms. **B**. Distribution of Ks between a pair of homologous genes between oil palm and date palm (speciation), and separately within oil palm and date palm (WGD). **C**. Pairwise transposon divergence throughout 30 randomly selected subfamilies of LTR retrotransposon superfamily Copia in oil palm, where two major peaks at an average similarity of 82% and 92% are revealed. **D**. Pairwise transposon divergence throughout 30 randomly selected subfamilies of LTR retrotransposon superfamily Copia in date palm, where two major peaks at an average similarity of 84% and 93% are only slightly visible. **E**. Comparison of the relative expression level between a pair of locally duplicated genes throughout 12 examined samples, where only one of each pair of homologous genes showing intact LTR insertion in gene feature. KD1.5, KD2.5 and KD3.5 indicate kernel samples at 1.5, 2.5 and 3.5 months after fertilization, respectively, while MD1.5, MD2.5, MD3.5, MD4.5 and MD5.5 indicate mesocarp samples at 1.5, 2.5, 3.5, 4.5 and 5.5 months after fertilization, respectively. *, ** and ns indicate *P* < 0.05, *P* < 0.01 and not significant for paired *t*-test, respectively.

We did not find notable evidence of gene loss in date palm in contrast to oil palm, leading to another hypothesis that transposon proliferation drives genome size expansion and speciation of palms. LTRs are the richest transposable elements (TEs) in both species, with a total length of ∼ 950 Mb and ∼ 200 Mb for oil palm and date palm, respectively (Table S6). Difference in LTRs content explains ∼ 80.3% of the genome size variation between the two species. Among LTRs, Copia is the most abundant superfamily for both species and accounts for ∼ 52.3 % of total genome size variation (Table S7). We examined the historical dynamics of each subfamily of Copia. Pairwise sequence divergence within each subfamily presented two peaks with sequence similarity of ∼ 82% and ∼ 92%, respectively, in oil palm (**Figure 2C**). In comparison, in date palm, two peaks of sequence divergence at ∼ 84% and ∼ 93 % were only slightly visible (**Figure 2D**). In particular, the peak of higher similarity was remarkably inflated in oil palm, suggesting recent transposon burst in oil palm, relative to date palm. We compared sequence divergence between Copia and homologous genes that mark the last WGD and diversification. Homologs in conserved syntenic blocks within each species presented a consistent peak with similarity of ∼ 86%, while divergence of homologs between species was ∼ 92%, marking the WGD and diversification events, respectively (Figure S7). The first wave of transposon burst, overlapping with the last WGD event, is suggested to be caused by rediploidization due to elevated genomic stress soon after WGD [16]. Under the assumption of comparable sequence evolutionary rates [16], the second wave of transposon burst coincides with the time of divergence between the two palms, suggesting that transposon burst and differential dynamics play an important role in diversification of palms. Large-scale transposon proliferation and movement may drive chromosome rearrangements, variation of recombination and gene diversification, and eventually lead to speciation [17].

Transposon dynamics can affect genome-wide expression patterns and promote divergence by epigenetic regulation [18]. We identified 9,786 intact LTRs throughout the genome. Approximately 33.8% of the intact LTRs showed an estimated insertion time of < 1 Mya (Figure S8), within which 21.8% was located in or within 5-kb distance to gene features (Figure S9). We hypothesized that young intact LTRs closely linked to genes are able to affect gene expression patterns. First, we compared the expression levels of 273 pairs of paralogous genes from conserved syntenic blocks of oil palm, where only one of a pair of genes is closely linked to a young intact LTR. However, the expression levels of genes linked to LTRs were only slightly reduced (but not significantly) in almost all examined 12 tissues (Table S8; Figure S10). These paralogs diverged since the last WGD at ∼ 75 Mya [15] and likely have functionally diverged in depth. Thus, the effects of LTRs on linked genes are likely underestimated in these anciently duplicated genes. As expected, these paralogous genes presented a more diverse expression pattern than the recent locally duplicated genes (Figure S11). We further examined the effects of intact LTRs on 103 pairs of locally duplicated genes, with a younger duplication time (Ks, 0.15 ± 0.07). Interestingly, genes linked to LTRs showed significantly lower expression levels, in comparison to their adjacent paralogs, in most of the examined samples (**Figure 2E**). Similar results were reported in the tea plant (*Camellia sinensis* var. *sinensis*) [19]. These findings highlight the important roles of LTRs in promoting transcriptional diversification of duplicated genes by epigenetic suppression of closely linked genes, which finally contributes to genome divergence.

### Convergent evolution of the *VIRESCENS* gene in palms

It has been hypothesized for decades that fruit color has evolved to increase visual conspicuousness, and is subjected to selection by seed dispersing animals [20]. In tropical palms, fruit color evolution is suggested to have interactions with frugivores [21]. However, the genomic basis of adaptive evolution of fruit color in palms is still unclear. *VIRESCENS*, a R2R3-MYB transcription factor controlling the accumulation of anthocyanins in fruit exocarp of palms, leads to deep violet to black fruit colors [22]. The dark pigments of the exocarp reduce the visual conspicuousness in contrast with red, orange and yellow pigmented fruits, which are caused by carotenoids and carotenes, making it less attractive to herbivorous animals. Thus, genes controlling the accumulation of anthocyanins in the exocarp, tend to be under selection by seed dispersing animals.

It has been shown that loss-of-function mutants of *VIRESCENS* in both oil palm and date palm is associated with loss of anthocyanins in exocarp, leading to conspicuousness of fruit colors [10, 22]. To examine the hypothesis, we cloned and analyzed the *VIRESCENS* gene of four additional palms: coconut, Christmas palm (*Adonidia merrillii*), Macarthur palm (*Ptychosperma macarthurii*), and golden cane palm (*Dypsis lutescens*). First, we measured the absorption spectrum of exocarp extracts and found that all four palms were deficient in anthocyanins accumulation in ripe fruit exocarp, in comparison to oil palm *VIRESCENS* fruit (wild type) which had a high concentration of anthocyanins (**Figure 3A** and **B**). All coconut assemblies including the Catigan Green Dwarf [23], Cn. tall and Cn. dwarf [24] were observed to harbor the complete *VIRESCENS* locus (**Figure 3C**). Interestingly, sequence analysis showed that the *VIRESCENS* gene in these genome sequences was consistently disrupted by an insertion of a highly repetitive region including ∼60 simple repeats and ∼50 LTRs, and partially lost exon 1 and the whole of exon 2 (**Figure 3C**). Disruption of *VIRESCENS* in the genome likely explains the green exocarp even in ripe fruit of coconut. In Macarthur palm, we found a premature termination codon in the exon 3 of *VIRESCENS*, resulting in a predicted truncation of 34 amino acids in the C-terminal relative to the sequence of wildtype oil palm (**Figure 3D**). As predicted in both oil palm and date palm, the truncated 34 amino acids are overlapping with the transcriptional activation domain of R2R3-MYB transcription factor (Figure S12) and loss of this domain leads to deficiency in regulation of anthocyanins accumulation [22]. In contrast, we did not identify evidence of loss-of-function mutations in the coding sequences of *VIRESCENS* for both Christmas palm and golden cane palm (Figure S12). However, the expression of this gene was undetectable in ripe fruit exocarp of both palms (**Figure 3E**), implying that sequence variations in the regulation regions have silenced *VIRESCENS* in this lineage. Taken together, our data address convergent evolution of *VIRESCENS* gene in palms, under the selective pressure of seed dispersing animals.

**Figure 3.**
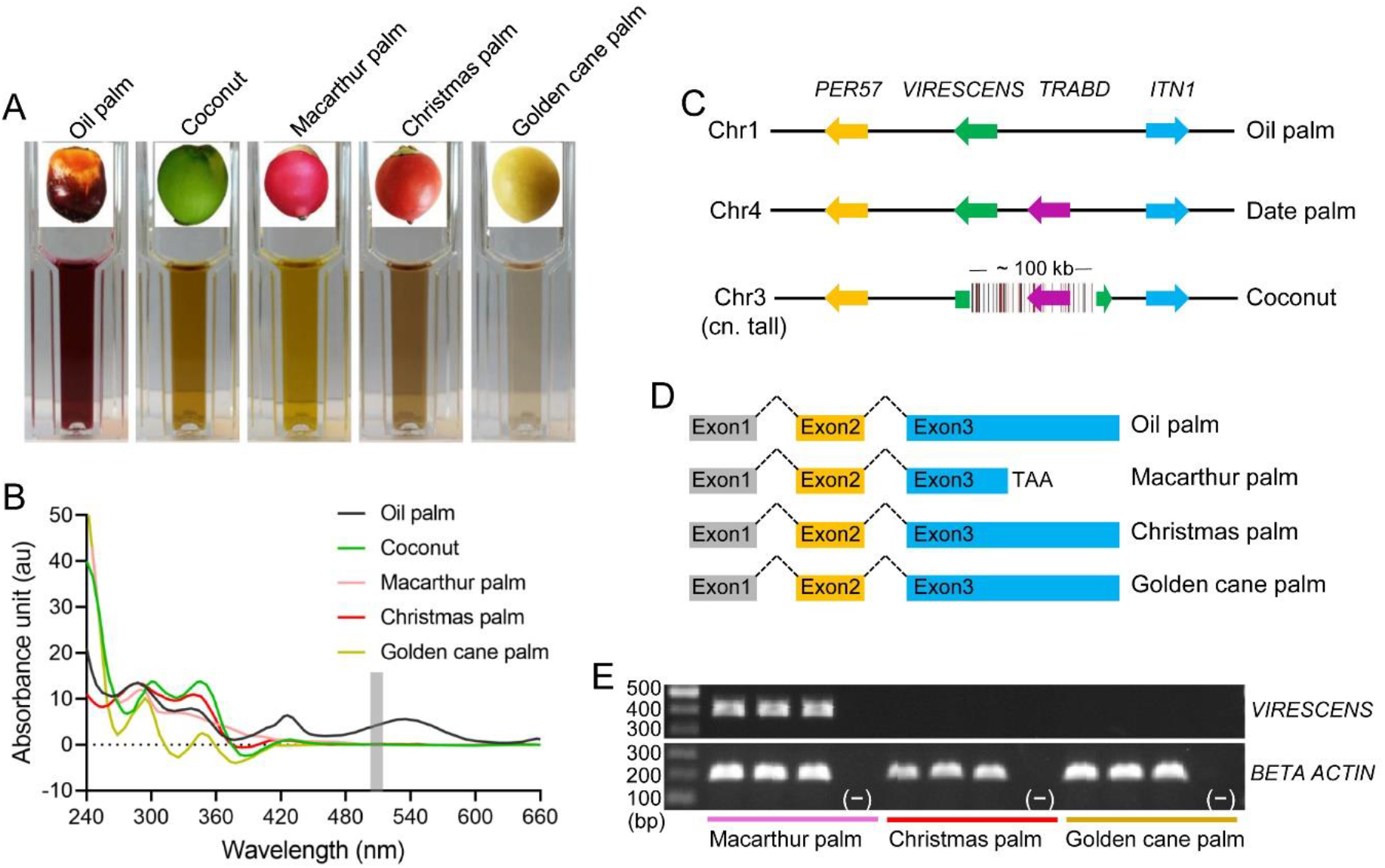
Variation of *VIRESCENS* gene in palms. **A**. Color of exocarp extracts in 1% acidified methanol across oil palm, coconut, Macarthur palm, Christmas palm and golden cane palm, where oil palm *VIRESCENS* fruit type containing anthocyanin, showed a dark purple color. **B**. UV absorption spectrum of exocarp extracts in 1% acidified methanol across palms, where oil palm *VIRESCENS* fruit type presented an absorbance peak at ∼ 530 nm, consistent with the absorption of anthocyanins. **C**. Genomic synteny of *VIRESCENS* locus among oil palm, date palm and coconut. Coconut *VIRESCENS* is disrupted by an insertion of 100-kb highly repetitive sequence, where black and red bards indicate simple repeats and LTR transposons, respectively. **D**. Gene models of *VIRESCENS* gene in oil palm, Macarthur palm, Christmas palm and golden cane palm, where a premature termination codon was detected in exon 3 of Macarthur palm. **E**. Expression of *VIRESCENS* gene and housekeeping gene (*BETA ACTIN*) in Macarthur palm, Christmas palm and golden cane palm, examined using gene-specific primers. Three individuals were examined for each species and minus indicates negative control.

### Duplication of PR genes and responses to *Ganoderma boninense* in oil palm

Pathogenesis-related proteins (PRs), subgrouped into functionally different groups in plants, play critical roles in host defense to viral and fungal infections [25]. To date, however, little is known about the mechanism of these proteins responding to pathogen infections. We discovered 505 PRs from 16 families in oil palm genome, among which 483 were mapped in 16 chromosomes (**Figure 4A**; Table S9). We found 319, 382 and 427 PRs in date palm, coconut and banana genome sequences, respectively. The size of gene families in oil palm was well correlated with that in date palm, coconut and banana (*R* > 0.92, *P* < 10^− 4^) (Figure S13), showing no significant evidence of expansion for a specific family. In oil palm, most of the PRs presented in tandem duplications (**Figure 4A**). We defined a tandem array as a region, within which genomic distance between any two adjacent PRs was < 100 kb. We discovered 70 tandem arrays, with the number of PRs within each ranging from 2 to 23. Over 64.4% (312) of PRs were found to be located in tandem arrays. The largest tandem array was located at chr1, consisting of 23 members of PR16 (**Figure 4A**). We observed that ∼ 97% of PRs in individual tandem arrays were resulting from tandem duplications, while the remaining ∼ 3% were from translocation or ancient duplication and divergence. Interestingly, we did not find obvious evidence that these tandem arrays were distributed between a pair of conserved syntenic chromosome blocks. These data suggest that PRs are hyperactive in birth and death, as well as translocations and may have frequently reorganized their genomic locations.

**Figure 4.**
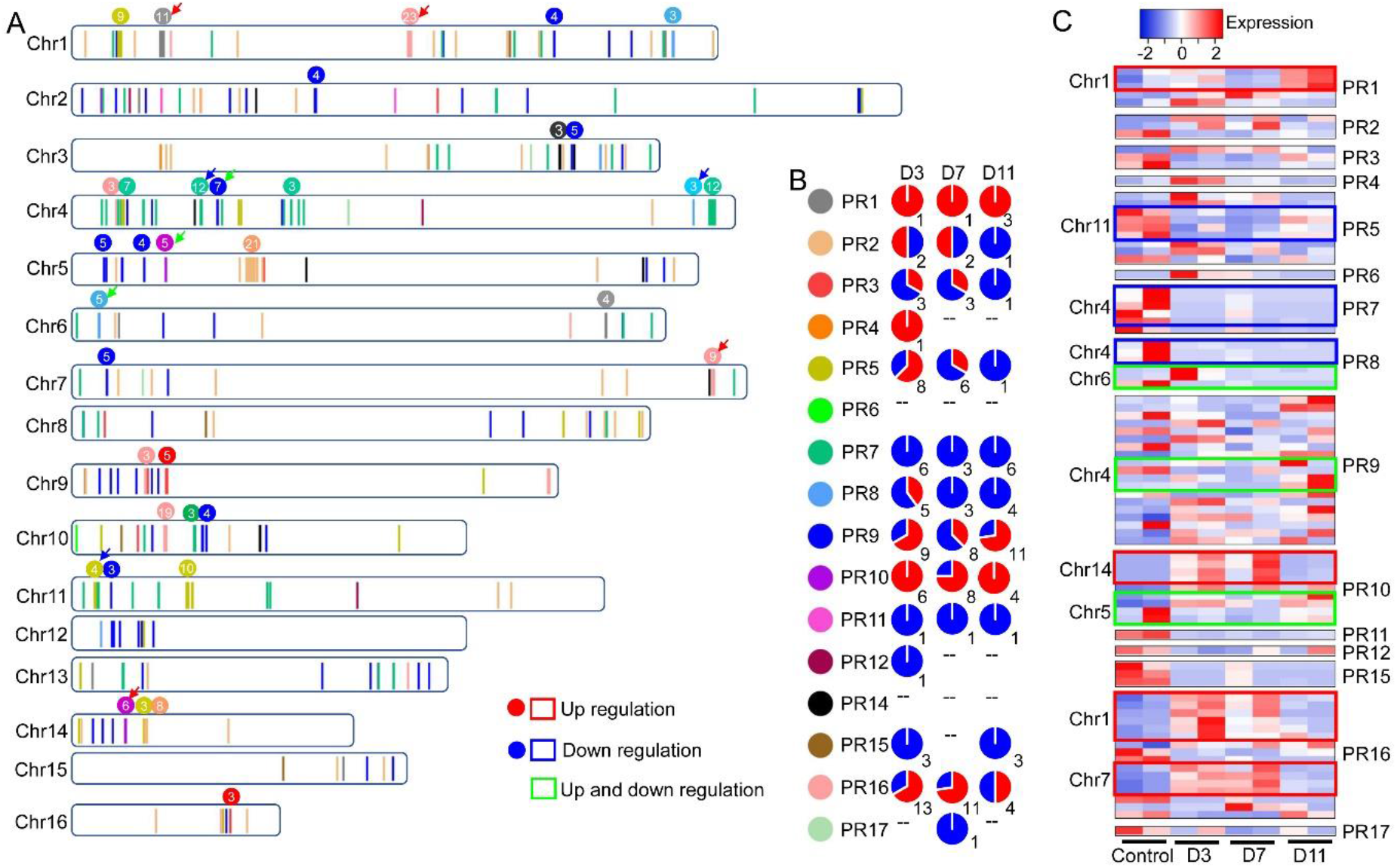
Genome-wide distribution and the relative expression of pathogenesis-related proteins (PRs) in oil palm. **A**. Genome-wide distribution of PRs throughout 16 chromosomes. Positions of PRs are shown with vertical bars and PR families are discriminated by colors as shown in **B**. The number and family of PRs in tandem arrays are indicated at the top of each array. Red and blue arrows indicate that PRs in the tandem arrays show consistently up and down regulation, respectively, while green arrow indicates PRs in the tandem arrays show both up and down regulation, against fungal infection. **B**. Pie chart shows the number of significantly up- and down-regulated PRs throughout 16 PR families, in oil palm root, at three, seven and 11 days post fungal infection. **C**. Heatmap showing the relative expression of PRs that are located in tandem arrays, in oil palm root, at three, seven and 11 days post fungal infection. Tandem arrays, within which PRs show consistently up and down regulation, are highlighted with red and blue boxes, respectively, while those show both up and down regulation are highlighted with green box.

To understand more about the mechanism of pathogen defense in oil palm, we analyzed the genome-wide expression pattern of PRs against the infection by *Ganoderma boninense* (Table S10) published by others [26]. We found that 84 (16.7%) PRs were among the reported differentially expressed genes (DEGs) in root transcriptomes post infection (Table S11; Figures S14 and S15). The remaining PRs may be induced in other tissues or involved in responses to the other pathogens. In detail, 59, 47 and 39 PRs were detected as DEGs at 3, 7 and 11 days post infection (dpi), respectively (Figures S14 and S15). Thirty-eight (45.2%) DEGs were located in 10 tandem arrays, and distributed across seven chromosomes: chr1, chr4, chr5, chr6, chr7, chr11 and chr14 (**Figure 4A**). DEGs in four (chr1:PR1s, chr1:PR16s, chr7:PR16s and chr14:PR10s) and three (chr4:PR7s, chr4:PR8s and chr11:PR5s) tandem arrays were consistently up and down regulated, respectively (**Figure 4A**). Analysis of DEGs in individual families (*e*.*g*., PRs in PR5, PR9 and PR16 families) did not always show a consistent expression pattern (**Figure 4B**), suggesting neofunctionalization of the differentially expressed PRs. Notably, DEGs belonging to PR16 family were largely located in two tandem arrays at chr1 and chr7, where all DEGs were up-regulated (**Figure 4C**), implying that these PRs are subfunctionalized and involved in additive resistance to *G. boninense* [27]. Phylogenetic analysis revealed that three major genetic clusters in PR16 family and the identified DEGs were all from the subclade 4, the youngest subclade of clade 3 (Figure S16). With regard to another tandem array at chr10, we found that PR16 members were from different subclades and showed a chimeric pattern of organization. In this tandem array, PRs of subclade 2 and 3 as a unit were repeatedly organized (Figure S17), as observed in some other plants, *i*.*e*., *Theobroma cacao* and *Manihot esculenta* [28, 29], suggesting that PRs have diverged prior to tandem duplications over evolutionary time. Taken together, our results revealed the crucial roles of large tandem arrays of PRs in defense responses, particularly those consisting of evolutionary closely related PR genes. PRs in chimeric tandem arrays or showing expression pattern shifts could have diverged over evolutionary time and likely been neofunctionalized and/or subfunctionalized.

### Population structure of the African oil palm

We first examined population structure of oil palms based on 4,410,076 SNPs generated by whole-genome re-sequencing of 72 trees (**Figure 5A**; Table S12). Both Principal Component (PCA) and Admixture analyses showed that the oil palms in Southeast Asia have clearly differentiated from their ancestral African ones except for those from Singapore and Malaysia, where most of the oil palms were either assigned into the African cluster or differentiated into an intermediate cluster between the African and Southeast Asian clusters (**Figure 5B** and **C**), since the first introduction at ∼ 1840s [4]. Within Africa, oil palms from the Ivory Coast are strikingly differentiated from the remaining trees. Oil palms from Ghana, Nigeria, Cameroon and Angola formed into the other cluster, where pairwise differentiation among locations is limited, with an overall *F*_*ST*_ of ∼ 0.05 (Table S13). The localized oil palms of Southeast Asia showed considerable differentiation among each other. Admixture suggested the most likely number of genetic clusters to be three, followed by two (Figure S18). In agreement with PCA, Admixture showed evidence of a mixture of genetic clusters between oil palms of Southeast Asia and Africa. The mixture occurred only in the oil palms of Singapore and Malaysia from Southeast Asia, implying repeated introduction of oil palms to Southeast Asia, likely as a result of escape of commercial cultivation or frequent commercial trade in the studied area.

**Figure 5.**
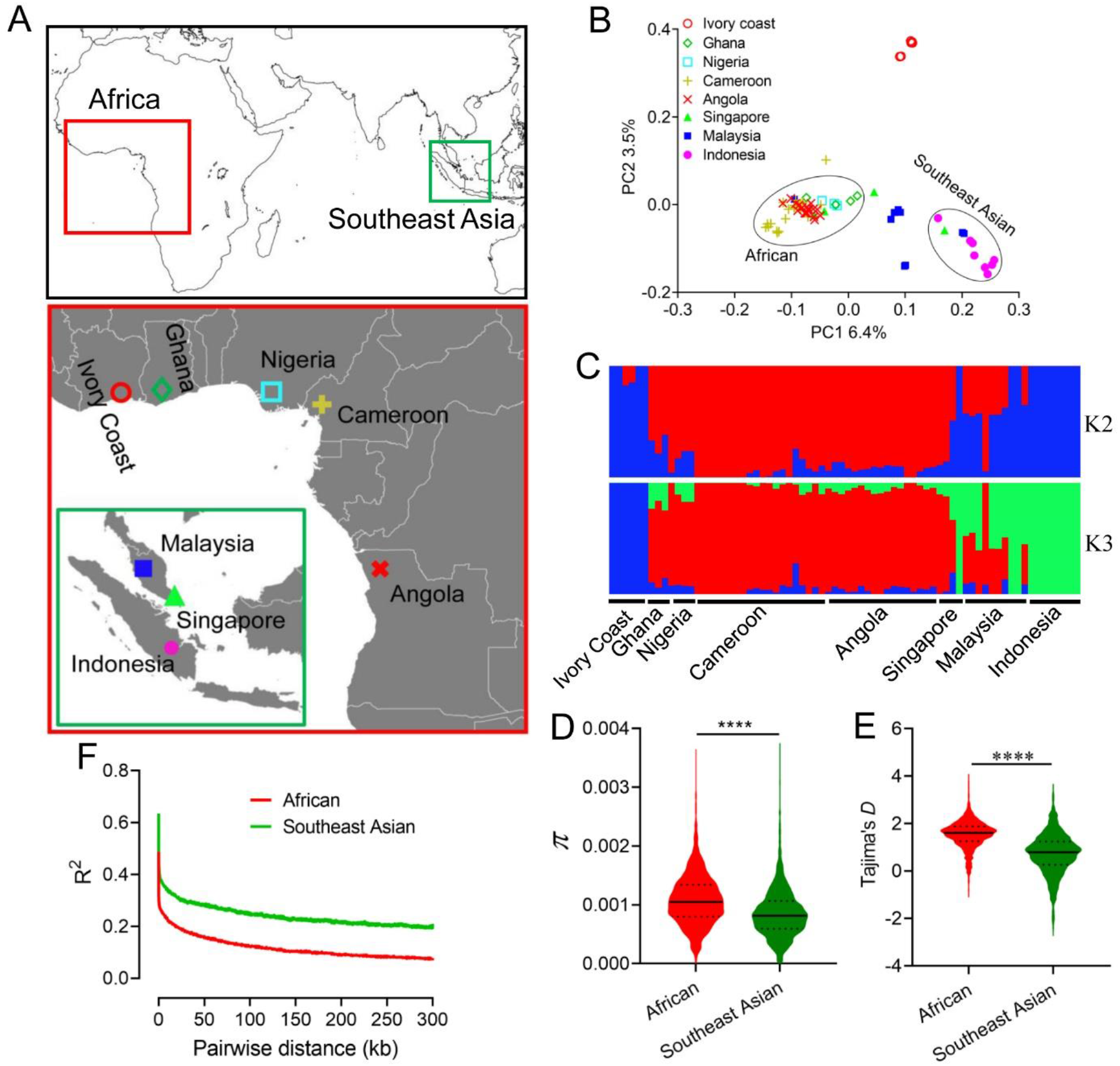
Population divergence of oil palm. **A**. Sampling localities of the ancestral African oil palms and localized Southeast Asian oil palms. **B**. Population structure between and within African and Southeast Asian oil palms, revealed by Principal Component Analysis. **C**. Genetic clusters of population ancestry between and within African and Southeast Asian oil palms, inferred with Admixture. **D**. and **E**. Comparison of nucleotide diversity (π) and Tajima’ *D* between African and Southeast Asian oil palms, estimated with 100-kb window size, where *P* value for *t*-test is shown above violin plot. **F**. LD decay separately in African and Southeast Asian oil palms.

Introduction of species would lead to loss of genetic diversity, as a result of founder effects and local selection in the new habitats. We examined the genetic diversity between African and Southeast Asian oil palms. Compared to their ancestral populations, oil palms in Southeast Asia have significantly reduced in genetic diversity, measured in nucleotide diversity (π: 0.0008 vs 0.0011, *P* < 10^− 48^, *t*− test) (**Figure 5D**), suggesting recent bottleneck and/or local selection during establishment of the Southeast Asian populations. We also observed significantly more negative Tajima’s *D* in the Southeast Asian oil palms, in comparison to the African oil palms (*P* < 10^− 72^, *t*− test) (**Figure 5E**), suggesting elevated positive selection in the localized Southeast Asian oil palms. We further estimated the linkage disequilibrium (LD) and found that LD decayed to half of the maximum within 10 kb in the oil palms of Africa, faster than that in Southeast Asia with a value of 30 kb (**Figure 5F**). This scale of LD allows effective identification of signature of selection using genome-wide SNPs.

### Adaptive evolution of the African oil palm

To identify signature of selection during introduction, we conducted whole-genome scan for candidate regions between African and Southeast Asian oil palms. Both PCA and Admixture showed that the 72 oil palms were split into two major genetic clusters (**Figure 5B** and **C**). *F*_ST_ and ϴ_π_ scans identified 127 consistent genomic regions under putative selection, with a total length of ∼ 23 Mb (1.3%) and containing 488 predicted protein coding genes (**Figure 6A** and **B**). Sixty-four out of the 127 regions deviated from neutrality by Tajima’s D analysis. A total of 317 genes were identified in those regions. Only the consistent results of these genomic scans were considered for further analysis to obtain a confident and reliable result (Table S14). Gene ontology enrichment analysis showed that these genes were more involved in stress responses, such as response to UV, regulation of autophagy and response to oxidative stress (Figure S19A). PFAM enrichment analysis revealed a notably large proportion of protein families were related to stress responses and disease defense, such as the GDA1/CD39 family, cytochrome P450, monooxygenase and haem peroxidase. In addition, genes from the ZIP Zinc transporter, sodium/hydrogen and transmembrane domain of ABC transporters protein families that are related to ion transport were also enriched (Figure S19B). Three genes (two homologs of *WRKY70* and *WRKY24*) from the transcription factor family WRKY were under putative selection. Genes of this family have been extensively shown to be related to abiotic stress in both model plants and oil palm [30]. These results suggest that genomic regions under putative selection play an important role in adaptive evolution of oil palm.

**Figure 6.**
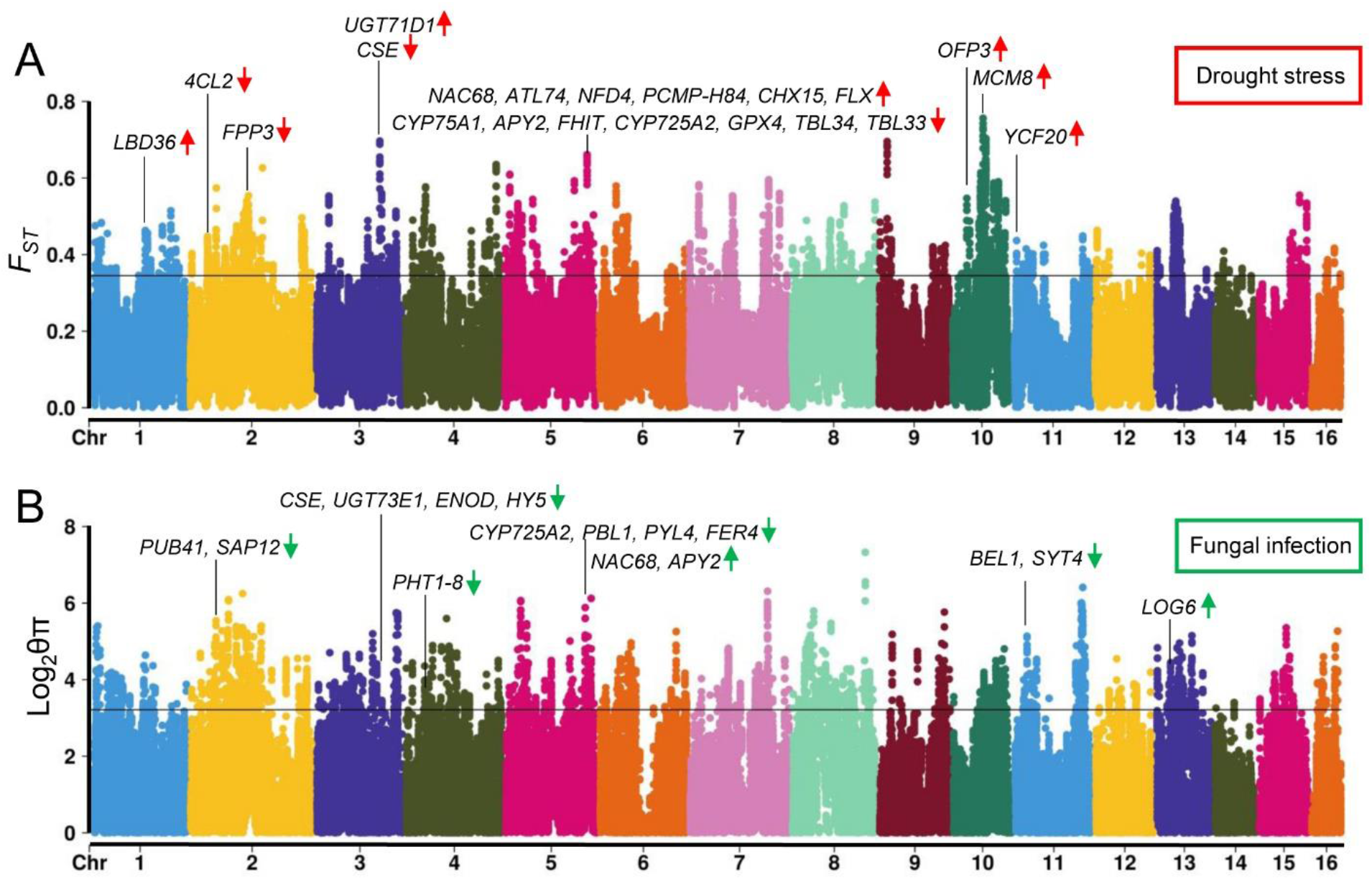
Signature of selection in oil palm. **A**. Manhattan plot of genomic regions under putative selection, revealed by *F*_*ST*_ scan between African and Southeast Asian oil palms. Genome-wide significance threshold at top 5% of windows in the empirical distribution is shown. Genes within outlier regions and identified as DEGs in root transcriptomes against drought stress are highlighted with gene names. Up and down oriented red arrows indicate up- and down-regulated DEGs, respectively. **B**. Manhattan plot of genomic regions under putative selection, revealed by *ϴ*_*π*_ scan between African and Southeast Asian oil palms. Genes within outlier regions and identified as DEGs in root transcriptomes against fungal infection, are highlighted.

As the genes under putative selection are more involved in stress responses, in particular, to pathogen infection and ion homeostasis, we separately analyzed these genes in correlation with DEGs responsible for resistance to *Ganoderma boninense* infection as described above and drought stress in our previous study [31]. Out of the 317 genes, 21 were revealed to be DEGs for drought stress, where 11 and 10 genes were up- and down-regulated, respectively (**Figure 6A**). Interestingly, we identified a selected region located at chr5: 105910001-110488835 with a length of ∼ 4.5 Mb, where 13 (14.3%) out of 91 genes were DEGs, significantly higher than the ratio under null hypothesis of 4.3% throughout the whole genome. Most of these genes have been verified to be responsible for drought resistance in model plants, such as *4CL2, CYP75A1, APY2, NAC68, CHX15, TBL33* and *TBL34* [32-35]. Some of these genes under putative selection were also revealed to be responsible for heat stress, like *LBD36, NAC68*, and *YCF20, 4CL2* and *CYP75A1* [36-38].

Among the putative selected genes, 16 were identified as DEGs against *G. boninense* infection, where 3 and 13 were up- and down-regulated, respectively (**Figure 6B**). Some of these genes have been shown to associate with disease resistance in model plant species, such as *BEL1, PUB41, SAP12, CSE, NAC6*8, *PBL1* and *PYL4* [33, 39-42]. Interestingly, most of these genes were down-regulated against infection, when up-regulation of these genes acts to enhance disease resistance [42, 43], implying a potential for decreased disease resistance in the oil palms of Southeast Asia. Three genes: *CSE, NAC68* and *CYP75A1* were observed to be responsible for both drought tolerance and fungal resistance, suggesting these genes play common roles in stress responses. Further functional studies of these genes would provide insights on adaptive evolution of oil palm out of Africa and also valuable resources for selective breeding of the species.

## Conclusions

We sequenced and assembled a chromosome-level genome sequence of the African oil palm. The genome assembly is of high completeness and continuity, and serves as a reference genome for oil palm. Comparative genomic analysis revealed that historical transposon expansion, but not WGD, explains genome size variation of palms and providing essential resources for adaptive radiation. The *VIRESCENS* gene, with its association with fruit color variation, is likely under selection during adaptive radiation of palms. Tandem highly repeated genes, PRs play an important role in defense responses to *Ganoderma* infection. Analysis of genetic variation between the ancestral African and recently introduced Southeast Asian oil palms identified signatures of selection, particularly on the introduced oil palms. Genes under putative selection are remarkably associated with stress responses, providing insights into adaptation to new habitats. The novel genomic resources and insights gained from this study could be exploited for comparative genomics, evolutionary studies and genetic improvement of palms.

## Materials and methods

### Genome sequencing and assembly in oil palm

The same *Dura* tree, previously sequenced with Illumina platform [6], was sequenced using SMRT technology to improve the genome assembly. Genomic DNA was isolated with MagAttract HMW DNA Kit (Cat. No. 67563, Qiagen, Germany). Two 20-kb libraries were constructed and sequenced for > 150× coverage on PacBio Sequel II (Pacific Biosciences, CA) by BGI (Hongkong, China). Flye v2.8 [44] was used to assemble the genome (-g 1.8g -m 10000 --asm-coverage 50 -i 3). Cleaned paired-end reads of 300-bp insert libraries and ∼ 100× coverage from Illumina sequencing [6] were used to polish the genome with Pilon [45].

### Construction of high-density linkage maps

For constructing high-density linkage maps, five F_2_ families consisting of a total of 978 progenies were used for RAD sequencing. DNA was isolated from leaves of each tree using DNeasy Plant Mini Kit (Cat. No. 69104, Qiagen). DNA was digested with PstI-HF restriction enzymes (R3140L, New England Biolabs, MA) and RAD libraries were constructed as described in our previous study [46]. The libraries were sequenced for 150 bp single-end reads on NextSeq500 platform (Illumina, CA). Raw reads were cleaned with process_radtags (-r -c -q -t 130) in Stacks package [47]. Cleaned reads of ∼ 7.3 million for each sample were aligned to the above reference genome with BWA-mem [48] with default parameters. Aligned reads were assembled and called for SNPs with Stacks package [47], according to our previous study [46]. Only one SNP from each RAD tag was kept. SNPs that were present in > 90% individuals within each family and showed Mendelian segregation distortion of > 0.05 in Chi-squared test were retained. Linkage mapping was carried out using Lep-MAP3 [49], with an LOD of > 10 for linkage group assignment.

### Construction of chromosomal-level genome assemblies of palms

RAD tags that were incorporated into the five high-density linkage maps were aligned to the contigs to assign genomic coordinates. Chimeric contigs were determined by linkage maps, which are not likely to have among-chromosome grouping errors [50]. Contigs with more than four markers mapped to different linkage groups, were considered as chimeric and were then split at the longest gaps between mismatched fragments. The program ALLMAPs [51] was then employed to anchor contigs to linkage maps, with default parameters. Centromere positions were estimated based on the distribution of recombination rates along individual chromosomes. Recombination rates, measured as ρ = 4Ner per kb, were estimated using LDhat [52]. Completeness of genome was examined by mapping to BUSCO v3.0.1 database [53].

### Repeat and genome annotation

RepeatModeler (http://www.repeatmasker.org) was first used to build a custom repeat library of the studied species. RepeatMasker [54] was then employed to identify repetitive sequences based on the custom repeat library and Repbase database [55]. Tandem repeats were further annotated using the program Tandem Repeat Finder [56]. Finally, we combined and filtered these repetitive sequences to obtain the nonredundant repeat annotations of the genome based on the coordinates. Assessment of the intact LTR retrotransposons (LTRs) was carried out using LTR_retriever [57]. Demographic history of the transposable elements (TEs) was inferred by investigation on the most abundant LTRs. One hundred LTRs were randomly selected from 40 random subfamilies of Copia. Full sequences were extracted and aligned with MUSCLE [58]. The distribution of pairwise sequence similarity within a family, was used to estimate the temporal dynamics of TE activity.

Genome was annotated with MAKER2 pipeline [59]. Genome sequences were first softmasked using RepeatMasker [54], based on the above repetitive libraries. Cleaned mRNA sequencing reads of multiple organs from our previous studies [6] were assembled with Trinity [60] and used for evidence-based annotation. For *ab initio* gene model prediction, protein sequences of *E. guineensis* EG5.1 [7] and EGv2 [6], date palm Barhee BC4 [10] and coconut HainanTall [61] were used as evidence. Programs SNAP [62] and Augustus [63] were iteratively used to train gene models. Predicated gene models that contain TE domains and are not supported by transcripts were filtered. Cleaned gene models were then annotated by blast to nr and RefSeq databases with BLASTP (E-value < 1e-10).

### Evolutionary analysis

Homologous blocks within and between species of interest were determined by pairwise whole-genome alignment with LASTZ [64] and all-versus-all blastp search with Ortholog-finder at gene level [65]. Putative one-to-one orthologs and paralogs from a pair of homologous blocks between oil palm and date palm and within oil palm, respectively, were aligned using MUSCLE [58]. Coding sequences were then aligned with the guidance of corresponding protein alignments. DNA alignments were further polished using trimAl [66]. Synonymous substitution rate (Ks) was estimated between a pair of homologous genes using KaKs_calculator [67]. Local duplicated genes were identified by analyzing the results of all-versus-all blastp search as described above, based on their genomic coordinates. To estimate the sequence divergence of LTR retrotransposons, 100 members from each of the 30 Copia subfamilies that were randomly selected, were randomly selected and pairwise aligned, according to a previous study [16]. Pairwise sequence divergence was estimated and compared to that of homologous genes to infer the relative evolution time [16].

### Transcriptome analysis

To compare the expression patterns of homologous genes, RNA sequencing reads of various organs from oil palm in our previous studies [31, 68, 69] and from date palm [70] were reanalyzed. Raw reads were cleaned with *process_shortreads* in Stacks package with default parameters, to remove adaptors and low-quality reads. Cleaned reads were then aligned to the reference genome using STAR [71], with default parameters. Uniquely mapped reads were counted to calculate gene expression level based on genome annotations, using the program HTSeq-count [72]. Gene expression level was then quantified as the number of fragments per kilobase of transcript per million mapped reads (FPKM). Heatmapper [73] was used to visualize the clusters and relative expression of genes.

### Analysis of *VIRESCENS* gene in palms

The presence of anthocyanins across palms was examined by measurement of the absorption spectrum of exocarp extracts in 1% acidified methanol, according to a previous method [22]. Equal exocarp material (100 mg) for each palm were used for extraction and the spectrum of UV absorption was measured from 240 nm to 700 nm with a 10 nm interval. Sequences of ***VIRESCENS*** gene across the studied palms were amplified either by amplification of genomic DNA or cDNA, using primers designed according to sequence homology among oil palm, date palm and coconut and primer walking (Table S15). Coding sequences were predicted based on oil palm ***VIRESCENS*** gene [22]. Predicted protein sequences were aligned using MUSCLE [58] and a phylogenetic tree was constructed using IQ-TREE2 [74], under HIVb+I model with 1,000 bootstrap replications. The relative expression of ***VIRESCENS*** gene was examined using reverse transcription PCR (RT-PCR). In brief, total RNA extraction and cDNA synthesis was carried out according to our previous study [31]. Quantity of cDNA corresponding to 50 ng of total RNA was used as template for amplification using gene-specific primers and the housekeeping gene, beta-actin, was used as a reference, with the following PCR condition: 94°C for 5min, followed by 35 cycles of 94°C for 30s, 60°C for 30s and 72°C for 30s, and a final extension of 72°C for 5min. RT-PCR products were examined by running 2% agarose gel.

### Characterization of PR proteins

Protein sequences of all PR family members of different plant species [28] were used as baits to search the genomes of oil palm, date palm, coconut and banana, with blastp (E-value < 1E-5). Protein sequences were extracted and manually curated, and were then sorted and classified based on protein domains, according to a previous study [28]. Genomic coordinates of PR genes in oil palm were extracted from annotation files to study the distribution and duplication patterns. Protein sequences of PR family members of interest were aligned using MUSCLE [58]. Alignments were refined using trimAl [66]. Phylogenetic trees were constructed using IQ-TREE2 [74], under automatically searched mutation model (JTT+R4).

Functions of PR proteins in disease resistance was studied by analyzing the RNAseq data set of oil palm seedlings infected with *G. boninense* inoculums at 3, 7 and 11 days post inoculation (dpi) [26]. Processing of raw sequencing reads, alignment to reference genome, count of mapped reads were carried out as described above. Normalization of transcripts and identification of differentially expressed genes (DEGs) were performed using DESeq2 [75]. Genes with a fold change of > 2 and a significant cutoff value of 0.005, corresponding to 0.1 after FDR corrections, were considered as DEGs.

### Whole-genome resequencing and variant calling

A total of 72 trees from West Africa (50) and Southeast Asia (22) were selected for sequencing. Libraries of 550 bp inserts were constructed using Truseq DNA PCR-Free kit and sequenced on NextSeq500 (Illumina, USA). Raw reads were filtered as described above. Cleaned reads were aligned against reference genome using BWA-mem [48] and variants were called using the Picard/GATK v4.0 best practices workflows [76]. We further filtered SNPs with the parameters: ‘QD < 2.0 ‖ DP > 5 ‖ FS > 60.0 ‖ MQ < 40.0 ‖ MQRankSum < − 12.5 ‖ ReadPosRankSum < − 8.0 ‖ SOR > 4.0’. Only SNPs were retained for further analysis and those with missing data across populations > 20 were also removed.

### Analyzing genetic diversity and population structure

Population genetic diversity including nucleotide diversity (π) and Tajima’s D, and pairwise differentiation (F_ST_) were estimated using VCFtools [77]. Population structure was analyzed with Principal Component Analysis (PCA) using Plink2 [78]. Genetic clusters from ancestry were inferred using Admixture [79], with the number of clusters ranging from 2 to 10. Cross-validation error was estimated to determine the most likely number of ancestral populations. Linkage disequilibrium (R^2^) between SNPs within populations was estimated using PopLDdecay (-MAF 0.02, -Het 0.88, -Miss 0.25) [80].

### Identifying signature of selection

Signature of selection between populations was inferred using F_ST_ and ϴπ (the ratio of nucleotide diversity between ancestral and introduced populations), and Tajima’s *D* statistics within Southeast Asian sample. These estimates were calculated in sliding window size of 100 kb with a window size step of 50 kb. Genomic regions consistently within top 5% of windows for *F*_*ST*_ and π, and bottom 5% of windows for Tajima’s D in the empirical distribution, were considered as outliers under putative selection. Protein coding genes in outlier regions were considered under putative selection. Protein sequences were extracted and annotated against Arabidopsis protein database (Ensembl TAIR10) using blastp with an E-value cutoff < 1E-10. Metascape [81] was employed to perform gene ontology enrichment analysis, with Arabidopsis as reference, using default parameters. DEGs responding to drought stress [31] and fungal infection as described above, were used to infer signature of selection under putative stresses.

## Data availability

Raw sequencing reads used in this study have been deposited to the DDBJ SRA database with an accession no. PRJDB11628. The chromosomal-level genome sequences of oil palm can be accessed from https://genhua.tll.org.sg/ and the China National GeneBank DataBase under Bioproject no. CNP0003017 (Assembly ID: CNA0047477).

## CRediT author statement

**Le Wang**: Methodology, Software, Formal analysis, Visualization, Writing – original draft preparation, Writing - review and editing. **May Lee**: Resources, Methodology, Formal analysis. **Zi Yi Wan**: Resources, Methodology, Formal analysis. **Bin Bai**: Resources, Methodology, Formal analysis. **Baoqing Ye**: Software, Formal analysis. **Yuzer Alfiko**: Resources, Methodology. **Ramadsyah Ramadsyah**: Resources, Methodology. **Sigit Purwantomo**: Resources, Methodology. **Zhoujun Song**: Resources, Formal analysis. **Antonius Suwanto**: Resources, Supervision. **Gen Hua Yue**: Conceptualization, Supervision, Resources, Funding acquisition, Writing - review & editing. All authors read and approved the final manuscript.

## Competing interests

The authors have declared no competing interests.

## Acknowledgements

This research was supported by Internal Funds of the Temasek Life Sciences Laboratory, Singapore (5020) and Wilmar International, Singapore (9200). We thank other lab members for technical supports.

## Supplementary material

**Figure S1 Distribution of SNPs in the five high-density linkage maps of the African oil palm**

**Figure S2 Anchoring contigs to five high-density linkage maps of the African oil palm, using the program ALLMAPS**

**Figure S3 Distribution of the annotation edit distance (AED) values throughout all predicted protein coding genes of the African oil palm genome**

**Figure S4 Comparison of expanded gene families between the African oil palm and date palm**

**Figure S5 Functional enrichment of the expanded gene families in the African oil palm, compared to date palm**

**Figure S6 Circos plot of homologous blocks between a pair of chromosome fragments in the African oil palm**

**Figure S7 Distribution of pairwise sequence divergence between homologous genes within oil palm and date palm (WGD event), and between oil palm and date palm (speciation event)**

**Figure S8 Distribution of estimated LTR insertion time, revealed by LTR_retriever**

**Figure S9 Distribution of the distance between intact LTRs and predicted protein coding genes**

**Figure S10 Comparison of the relative expression level between a pair of homologous genes from a pair of conserved chromosome syntenic blocks across 12 examined samples**

**Figure S11 Heatmap shows (upper figure) the relative expression level between a pair of homologous genes from a pair of conserved chromosome syntenic blocks across 12 examined samples**

**Figure S12 Alignment of virescens protein sequences and phylogenetic tree based on aligned protein sequences across oil palm, date palm, Macarthur palm, Christmas palm and golden cane palm**

**Figure S13 Correlation of the number of PR family members between the African oil palm and date palm, between the African oil palm and coconut, and between the African oil palm and banana**

**Figure S14 Number of differentially expressed genes (DEGs) identified in the root samples at 3, 7 and 11 days post infection with *Ganoderma boninense* (left) and the number of PRs that are identified as DEGs in these samples (right)**

**Figure S15 Total differentially expressed genes (DEGs) identified in the root samples at 3, 7 and 11 days post infection with *Ganoderma boninense* (green dots) and PRs that are identified as DEGs in these samples (red dots)**

**Figure S16 Phylogenetic tree constructed based on PR16 members and the relative expression of the PRs classified into the subclade 4, where red shade indicates consistently up-regulated genes in all infected samples, regardless of statistical power, while plus indicates statistically significant DEGs. ns, indicates non up-regulated genes**

**Figure S17 Genomic organization of PR16 members from different genetic clusters as revealed in Figure S18, in three tandem arrays at chr1, chr7 and chr10, respectively**

**Figure S18 Inferring the most likely number of genetic clusters (K) among African and Southeast Asian oil palms, based on cross-validation error**

**Figure S19 Gene ontology (A) and Pfam enrichments (B) of the genes under putative selection, consistently identified by *F***_***ST***_, ***ϴ***_***π***_ **and Tajima’ D scans, respectively**

**Table S1 Summary statistics of genome assemblies of oil palm, date palm and coconut**

**Table S2 Summary statistics of five high-density linkage maps, including the number of SNPs and length of each linkage group (LG)**

**Table S3 Summary statistics of SNPs that are mapped to contigs and genomic sequences that are anchored onto the linkage maps and oriented**

**Table S4 Genomic sequences anchored onto 16 linkage groups/pseudochromosomes of the African oil palm genome**

**Table S5 Completeness of oil genome assemblies assessed by BUSCO**

**Table S6 Summary statistics of transposable elements (TEs) in the African oil palm genome**

**Table S7 Summary statistics of LTR retrotransposon superfamilies in the African oil palm genome**

**Table S8 RNA sequencing samples used to examine homologous gene expressions in African oil palm genome**

**Table S9 Summary statistics of pathogenesis-related proteins (PRs) identified in the genome of oil palm, date palm, coconut and banana**

**Table S10 Root samples of African oil palm at 3, 7 and 11 days post infection with *Ganoderma boninense* and negative controls**

**Table S11 Differentially expressed PRs from different families at 3, 7 and 11 days post infection with *Ganoderma boninense***

**Table S12 African oil palm trees used for whole-genome resequencing and the sequencing coverage, sample location and date for each tree**

**Table S13 Pairwise F**_**ST**_ **of seven African oil palm samples, based on whole genome resequencing**

**Table S14 Genes identified under putative selection in the oil palm genome**

**Table S15 Genes that were identified under putative selection in the genome and to be differentially expressed in transcriptomes of oil palm against drought stress or fungal infection**.

**Table S16 Primers used to amplify and examine the expression of *VIRESCENS* gene in palms**

